# On the sensitivity of qPCR diagnostics for the canola clubroot pathogen *Plasmodiophora brassicae*

**DOI:** 10.64898/2026.02.05.704046

**Authors:** Heting Fu, Yalong Yang, Shiming Xue, Kher Zahr, Junye Jiang, Ronald Nyandoro, Jessica Haenni, Tiesen Cao, Michael W. Harding, David Feindel, Jie Feng

## Abstract

Clubroot, caused by *Plasmodiophora brassicae*, is an important disease of canola and other *Brassica* crops. Polymerase chain reaction (PCR), particularly probe-based quantitative PCR (qPCR), is widely used for the detection of *P. brassicae* in soil samples. To improve consistency in clubroot detection while maintaining efficiency, diagnostic laboratories would benefit from adopting a single, highly efficient qPCR system for routine testing. In this study, we analyzed the primer and probe sequences of all published PCR and qPCR systems for *P. brassicae* detection. Based on these analyses, three independently developed probe-based qPCR systems were selected and their performance was evaluated using synthesized target DNA (gBlock). One probe-based qPCR system exhibiting superior sensitivity on gBlock was subsequently evaluated on *P. brassicae* genomic DNA. This system consistently detected DNA equivalent to four resting spores per reaction, corresponding to a soil sample containing 1,000 spores per g soil when the DNA extraction protocol was considered as a component of the qPCR system. The sensitivity of the system was further validated using DNA extracted from soil samples collected from multiple locations across Alberta, where *P. brassicae* was detected at levels below those associated with visible clubroot symptoms. Based on these results, we recommend this qPCR system for routine clubroot diagnostics in laboratories across Canada.

## Introduction

Clubroot, caused by the protist *Plasmodiophora brassicae* Woronin, poses a major threat to Canadian canola (*Brassica napus*) production (Hwang et al. 2012). In the Canadian Prairies, clubroot was first detected on canola in 2003 in approximately a dozen fields near Edmonton, Alberta (Tewari et al. 2005). Since then, the disease has spread to over 4,000 fields in 47 Alberta’s counties and municipal districts (Strelkov et al. 2025) and has been confirmed in canola fields in Saskatchewan (Ziesman et al. 2019), Manitoba (Froese et al. 2019), Ontario (Al-Daoud et al. 2018), and North Dakota (Chittem et al. 2014).

*Plasmodiophora brassicae* produces large numbers of resting spores that can survive in soil for up to 20 years (Wallenhammar 1996). These resting spores serve as the primary source of inoculum under natural conditions. Consequently, both direct and indirect measurements of resting spores in soil have been widely used for the diagnosis of clubroot and forecasting clubroot risk. Numerous methods have been developed to detect resting spores in soil, among which PCR-based assays, including conventional end-point PCR, quantitative PCR (qPCR), and digital PCR, are considered the most sensitive and accurate (Faggian and Strelkov 2009; Wen et al. 2020).

The Alberta Plant Health Lab is responsible for clubroot evaluation including detection and quantification of *P. brassicae* resting spores in soil samples collected from canola fields across Alberta. These samples originate from provincial annual disease surveys as well as research activities focused on resistance breeding and the development of management strategies. Communication and collaboration with other diagnostic laboratories indicated that qPCR is the primary technique used for clubroot evaluation across Canada, with probe-based qPCR being more commonly employed than SYBR Green-based qPCR. However, considerable variation exists among laboratories with respect to the primers and probes used, and new primer and probe sets continue to be developed. While this expands the available options, it also creates uncertainty regarding which qPCR system provides the highest efficiency and reliability. To date, no comparative data were available to identify the most effective qPCR system for clubroot diagnostics.

In this study, we reviewed all published PCR and qPCR systems for *P. brassicae* detection and conducted a comparative evaluation of three independently developed probe-based qPCR systems. Our results identified one qPCR system with superior performance, and we recommend its adoption for routine use in diagnostic laboratories.

## Materials and Methods

### Chemicals and standard techniques

All chemicals and instruments were purchased from Fisher Scientific Canada (Ottawa, ON), unless otherwise specified. Primers, probes, and a gBlock were synthesized by Integrated DNA Technologies (Coralville, IA). DNA was extracted using the DNeasy PowerSoil Pro Kit (Qiagen Canada, Toronto, ON) on a QIAcube Connect instrument (Qiagen Canada). Extracted DNA was eluted in 50 µL of nuclease-free water. When required, DNA concentration was measured using a NanoDrop 1000 spectrophotometer.

### Primer sequence analysis

Sequences of all published primer sets used for PCR or qPCR detection of *P. brassicae* were retrieved (Fig. 1). The target regions of the primers were obtained by BLAST searching the primer sequences against the National Center for Biotechnology Information (NCBI) nucleotide collection (nr/nt) database (https://blast.ncbi.nlm.nih.gov/Blast.cgi). Primer sets developed for the detection of specific pathotypes (e.g., pathotype 5) were excluded. The remaining primer sets were grouped into three categories: conventional PCR, SYBR Green-based qPCR, and probe-based qPCR. Given the generally higher sensitivity of qPCR compared with conventional PCR, and the greater specificity of probe-based qPCR relative to the other two approaches, only probe-based qPCR primer sets were selected for further analysis. Primer dimer formation was predicted using the online Multiple Primer Analyzer (Thermo Fisher, https://www.thermofisher.com). Specificity of the primers in each qPCR system was confirmed by two BLAST analyses using the primer sequences as the queries: one against the NCBI nr/nt database and the other against the NCBI whole-genome shotgun (WGS) contigs database, with *P. brassicae* (taxid: 37360) excluded.

**Fig. 1.**
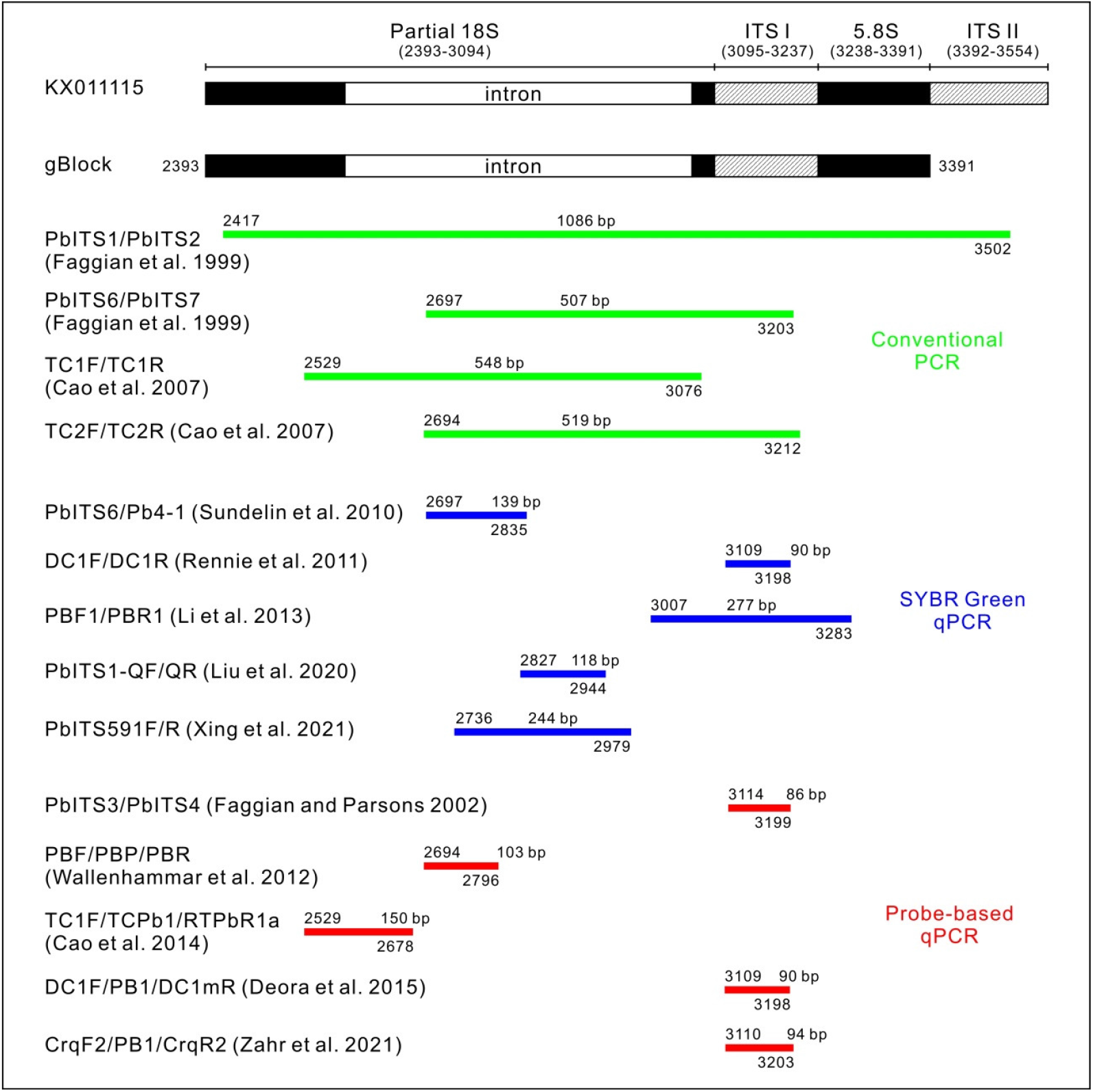
Diagrammatic sketch showing the genomic locations of targets of PCR and qPCR systems used for detection of the clubroot pathogen *Plasmodiophora brassicae*. KX01115 is the GenBank accession number for a partial ribosomal DNA (rDNA) sequence from a *P. brassicae* strain (https://www.ncbi.nlm.nih.gov/nuccore/KX011115). gBlock is a synthesized double-stranded DNA fragment used to evaluate the sensitivity of three probe-based qPCR systems. Numbers flanking each region indicate the start and end nucleotide positions within the KX01115 sequence.

### Probe-based qPCR reaction

Probe-based qPCR were conducted in PrimeTime gene expression master mix (Integrated DNA Technologies). Each reaction was in 20-µL volume containing 2 µL template regardless of the DNA concentrations, 0.25 µM of each primer, and 0.15 µM of the probe. The qPCR program consisted of an initial denaturation step at 95°C for 2 min, followed by 40 cycles of 5 s at 95°C and 30 s at 60°C. Each qPCR was conducted with three technical replicates.

### Sensitivity test of three probe-based qPCR systems on gBlock

A 999-base pair (bp) double-stranded DNA fragment was synthesized as a gBlock (Fig. 1). The gBlock sequence corresponds to nucleotides (nt) 2393-3391 of GenBank accession number KX01115, which represents a partial rDNA sequence of a *P. brassicae* strain. A series of 10-fold serial dilutions of the gBlock was prepared, ranging from 5 × 10^6^ to 0.5 molecules per µL. In addition, solutions at 1 and 2 molecules per µL were also prepared. Using the gBlock dilutions as templates, three probe-based qPCR systems developed by Wallenhammar et al. (2012), Cao et al. (2014), and Zahr et al. (2021) were tested. The experiment was repeated using an independent preparation of gBlock dilutions, yielding similar results and same conclusions.

### Sensitivity test of a probe-based qPCR system on *P. brassicae* genomic DNA

Clubroot galls on canola cultivar Westar were collected from a field nursery (53.6435, - 113.3493) at the Alberta Plant Health Lab and frozen at −20°C for three days. A resting spore suspension was prepared from the frozen galls following the method of Zahr et al. (2021). Using a haemocytometer, the suspension was serially diluted to a final concentration of 2.5-5 × 10^6^ spores per mL, corresponding to 10-20 spores per 0.04 mm^2^ square on the haemocytometer. The exact spore concentration was then determined as the mean of independent counts by three laboratory members, repeated until the standard deviation was less than one-tenth of the mean. A series of 10-fold serial dilutions of the resting spores was prepared, ranging from 1 × 10^9^ to 1 × 10^1^ spores per mL, and 100 µL of each dilution was subjected to DNA extraction.

A soil sample (black chernozemic; Pennock et al. 2011) was collected from a research field (53.6387, −113.3683) at the Alberta Plant Health Lab, autoclaved twice at 121°C, 15 psi for 1 h, and dried at 50°C for three days. The dried soil was ground with a coffee grinder and aliquoted into 1.5-mL microcentrifuge tubes at 100 mg per tube. Ten microliters of each resting spore dilution was added to the tubes, and the spore-soil mixtures were subjected to DNA extraction. Using the DNA from spore suspensions and spore-soil mixtures as templates, the probe-based qPCR system developed by Wallenhammar et al. (2012) was tested. The experiment was repeated using an independent preparation of spore dilutions and spore-soil mixtures, yielding similar results and same conclusions.

### Sensitivity test of a probe-based qPCR system on field soil samples

Soil samples of approximately 2 kg were collected from one field in each of nine counties in Alberta, all of which were found to have clubroot in previous surveys. Each soil sample was homogenized by hand shaking in a plastic bag, and a subsample (approximately 1 g) was taken. The subsamples were air-dried at room temperature overnight, and 100 mg of each was subjected to DNA extraction. Using the resulting DNA samples as templates, the probe-based qPCR developed by Wallenhammar et al. (2012) was tested. The experiment was repeated using an alternative preparation of subsamples from the same field samples, yielding similar results and same conclusions.

### Greenhouse assay on field soil samples

The remaining portion of each original soil sample was transferred to a 1-L square pot, into which ten seedlings of canola cultivar Westar were transplanted. The pots were maintained in a greenhouse. Seedling preparation and plant care, including temperature and light conditions, and watering, followed the protocol of Zahr et al. (2021). Six weeks after planting, the plants were removed from the pots and the presence of clubroot galls was visually checked. The experiment was not repeated.

### Data analysis

In all qPCR, means of the quantification cycle (Cq) values from the three technical replicates were calculated and treated as one data point. The qPCR standard curves were constructed by regression analysis using the SAS software (version 9.4; SAS Institute, Cary, NC). The efficiencies of qPCR primer sets were calculated as E = −1+10^(-1/slope)^ (Svec et al. 2015).

## Results

### Sequence analysis of published primers

To date, four conventional PCR, five SYBR Green-based qPCR and five probe-based qPCR systems were available for clubroot detection (Fig. 1). All of these systems target the rDNA region of *P. brassicae*. The sequences of primers and probes of these systems can be mapped to a 1086-bp region of the rDNA, for example, nt 2417-3502 of GenBank accession number KX01115 (Fig. 1). Analysis of primers from the five probe-based qPCR systems for potential primer dimer formation predicted self-dimerization for primer PbITS4 in the system developed by Faggian and Parsons (2002) and for primer DC1mR in the system described by Deora et al. (2015). These two systems were therefore excluded from further evaluation. BLAST analysis of primers and probes from the remaining three probe-based qPCR systems did not identify identical sequences in non-*P. brassicae* organisms, indicating that all three systems exhibit equivalent specificity for *P. brassicae* detection.

### Sensitivity test of three probe-based qPCR systems on gBlock DNA

The three probe-based qPCR systems developed by Wallenhammar et al. (2012), Cao et al. 2014, and Zahr et al. (2021) were tested on serial dilutions of gBlock DNA. A standard curve for each qPCR system was constructed based on Cq values obtained from these dilutions (Fig. 2). The calculated qPCR efficiencies of the three systems were 0.99 (Fig. 2a), 0.96 (Fig. 2b), and 1.00 (Fig. 2c), respectively.

**Fig. 2.**
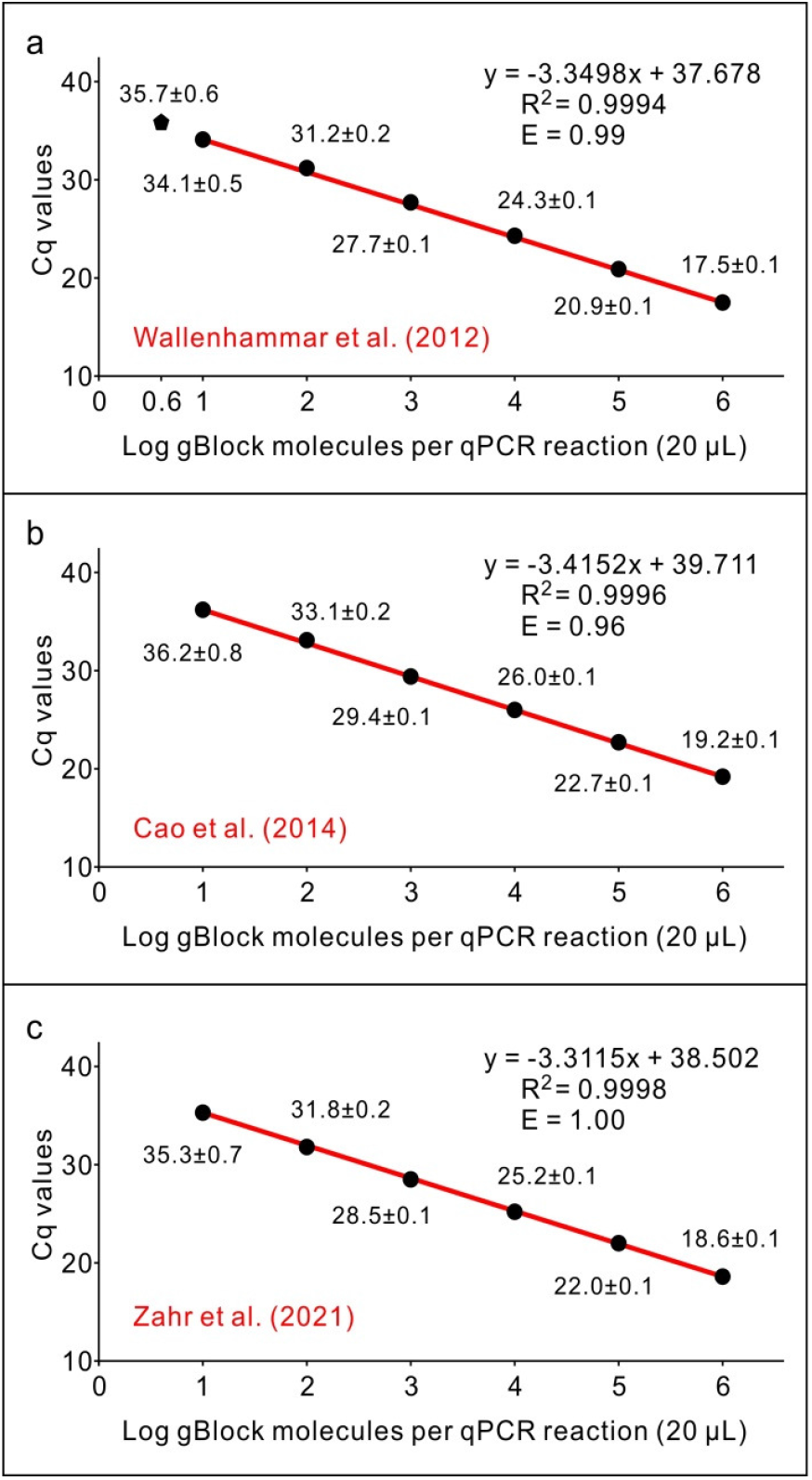
Sensitivities of three probe-based qPCR systems on gBlock DNA. a, the system developed by Wallenhammar et al. (2012). b, the system developed by Cao et al. (2014). c, the system described by Zahr et al. (2021). The qPCR standard curves were generated from the mean of quantification cycle (Cq) values against log10 of gBlock DNA copies in one reaction. The R^2^ score of the equation and the efficiency of the primers (E) are indicated over the curve. Efficiency was calculated as E = −1+10^(-1/slope)^. Each data point is shown as mean of three technical replicates ± standard deviation.

At equivalent gBlock DNA concentrations, the system developed by Wallenhammar et al. (2012) consistently produced lower Cq values than the other two systems. None of the three systems generated detectable amplification signals from 1 or 2 gBlock molecules (2 µL of 0.5 or 1 gBlock molecules per µL dilution was used in one reaction). However, the Wallenhammar et al. (2012) system generated a Cq value of 35.7 ± 0.6 from 4 gBlock molecules (Fig. 2a), whereas the other two systems failed to amplify at this concentration.

Based on these results, the Wallenhammar et al. (2012) system was identified as the most sensitive, with a detection limit of eight target copies per 20-µL reaction (4 molecules × two DNA strands per molecule).

### Sensitivity of the probe-based qPCR system on *P. brassicae* DNA

The Wallenhammar et al. (2012) qPCR system was further evaluated using DNA extracted from serial dilutions of *P. brassicae* resting spores and from spore-soil mixtures. Amplification signals were obtained from all tested samples (Fig. 3). The calculated qPCR efficiencies were 0.99 for spore-derived DNA (Fig. 3a) and 0.87 for spore-soil mixture DNA (Fig. 3b).

**Fig. 3.**
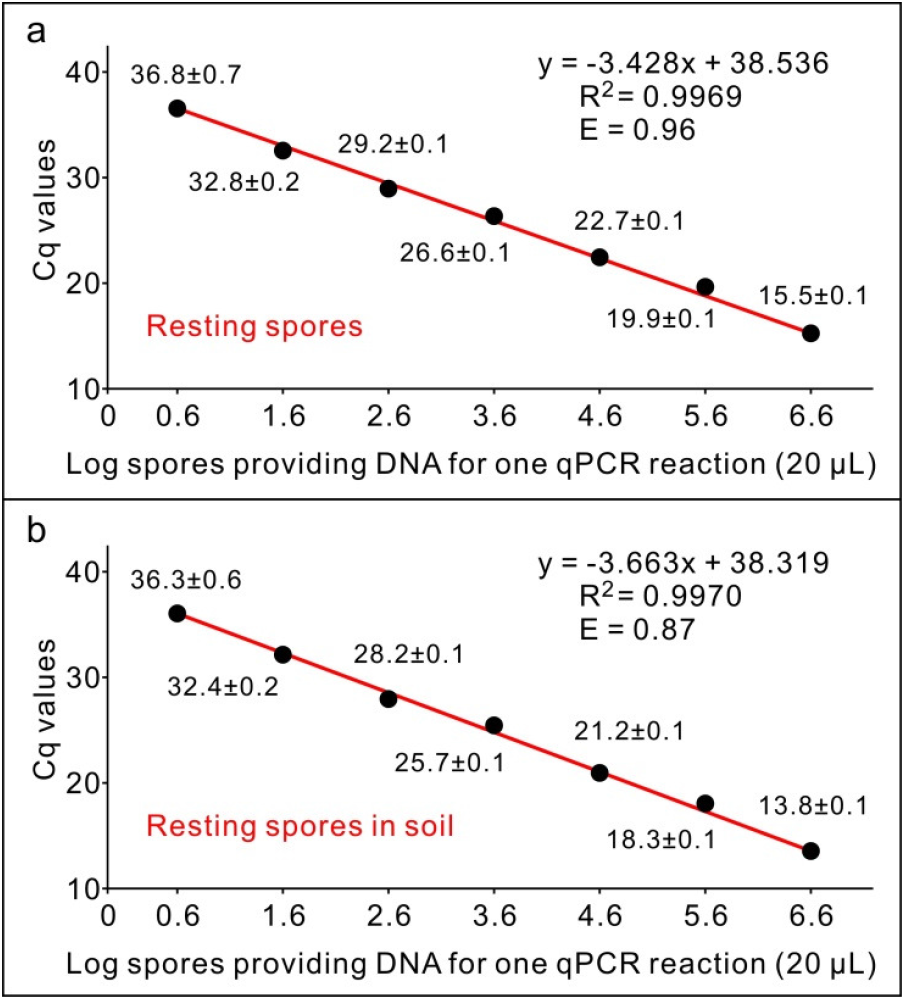
Sensitivities of the probe-based qPCR systems developed by Wallenhammar et al. (2012) on *Plasmodiophora brassicae* genomic DNA. a, DNA from serial dilutions of resting spores. b, DNA from mixtures of serial dilutions of resting spores and soil. The qPCR standard curves were generated from the mean of quantification cycle (Cq) values against log10 of spore number proving the template in one reaction. The R^2^ score of the equation and the efficiency of the primers (E) are indicated over the curve. Efficiency was calculated as E = −1+10^(-1/slope)^. Each data point is shown as mean of three technical replicates ± standard deviation.

For DNA derived from 100 resting spores and from a mixture of 100 spores-100 mg soil, mean Cq values of 36.8 ± 0.7 and 36.3 ± 0.6, respectively, were obtained. These results indicate that the detection limit from both spore DNA and spore-soil mixture DNA in a 20-µL reaction corresponds to DNA from approximately four resting spores. This estimate was calculated assuming the DNA was dissolved in 50 µL of water, and 2 µL was used as template per reaction. Since the sample was a mixture of 100 spores-100 mg soil, it is equivalent to a soil sample with a spore concentration of 1 × 10^3^ spores per g soil.

It has not escaped our notice that the Cq values obtained from spore DNA alone (Fig. 2a) were higher than those obtained from spore-soil mixture DNA (Fig. 2b) when equivalent numbers of spores were analyzed. This difference is likely due to the fact that the soil particles facilitate cell disruption during the initial DNA extraction step and thereby enhance DNA recovery. In addition, reduced qPCR efficiency (0.87) was observed when DNA templates were derived from serial dilutions of spores in soil (Fig. 2b). This effect is also likely related to the influence of soil particles, as higher spore concentrations would result in greater particle-mediated disruption of resting spores, leading to disproportionally increased DNA yields at higher spore concentrations.

### Sensitivity of the probe-based qPCR system on DNA from field soil samples

The Wallenhammar et al. (2012) qPCR system amplified signals from seven of the nine field soil samples. The corresponding greenhouse bioassay produced clubroot galls from six of these samples (Table 1). Comparison of qPCR and greenhouse assay results showed that soil samples with Cq values ≤ 34.4 consistently produced visible clubroot galls, whereas samples with Cq values ≥ 35.8 did not (Table 1). These results indicate that, under the conditions used in this study, the qPCR assay can detect *P. brassicae* resting spore levels below the threshold required to cause visually detectable root gall symptoms, confirming the high sensitivity of this qPCR system for clubroot detection.

**Table 1.** Sensitivity test of the probe-based qPCR system on soil samples.

| Sample Origin | Cq (Mean±SD) | Galls<sup>b</sup> |
| --- | --- | --- |
| 1 × 10<sup>5</sup> spores<sup>a</sup> | 26.5±0.1 | Y |
| Camrose County | 19.3±0.1 | Y |
| Leduc County | 26.3±0.1 | Y |
| Strathcona County | 27.1±0.1 | Y |
| County of Minburn | 28.9±0.1 | Y |
| Westlock County | 31.7±0.1 | Y |
| Sturgeon County | 34.4±0.4 | Y |
| Athabasca County | 35.8±0.4 | N |
| Lamont County | No signal | N |
| County of Vermilion River | No signal | N |
| Clean soil<sup>c</sup> | No signal | N |

<sup>a</sup>DNA was extracted from 1 × 10<sup>5</sup> spores, dissolved in 50 µL water and 2 µL was used in each qPCR reaction. Thus each reaction contained DNA from 4 × 10<sup>3</sup> spores.
<sup>b</sup>Presence (Y) or absence (N) of clubroot galls by visual detection.
<sup>c</sup>100-mg subsample from a soil sample that had been autoclaved twice at 121°C, 15 psi for 1 h, and dried at 50°C for three days.

## Discussion

The first published PCR system for clubroot detection was reported by Faggian et al. (1999). The first system in Canada was developed by Cao et al. (2007) and has been routinely used in diagnostic laboratories since its publication. The first probe-based qPCR system was published by Faggian and Parsons (2002). Targeting a rDNA region almost identical to that of Faggian and Parsons (2002), Rennie et al. (2011) published a pair of SYBR Green-based qPCR primers that differed by a few nucleotides from those reported by Faggian and Parsons (2002). Deora et al. (2015) later adopted the Rennie et al. (2011) primers, removing one nucleotide from the 3′ end of the reverse primer, and combined them with a probe identical to that of Faggian and Parsons (2002) to develop an alternative probe-based qPCR system. The reverse primers used in both Faggian and Parsons (2002) and Deora et al. (2015) were predicted to form self-dimers. To address this issue, Zahr et al. (2021) redesigned both the forward and reverse primers while retaining the probe from Faggian and Parsons (2002) and Deora et al. (2015). Because the systems described by Faggian and Parsons (2002), Deora et al. (2015), and Zahr et al. (2021) target nearly identical genomic regions and use identical probe, they were assumed to have comparable specificity. Primer dimer formation competes with target amplification for primer molecules and, more importantly, increases the accumulation of double-stranded DNA, which is a major factor inhibiting polymerase activity. Consequently, the Zahr et al. (2021) system was selected from these three for further evaluation. Among all five probe-based qPCR systems, those described by Wallenhammar et al. (2012) and Cao et al. (2014) were independently developed and target different rDNA regions. Therefore, these two assays were also selected for further study.

The reported detection limits of conventional PCR systems for *P. brassicae* are approximately 100 fg of genomic DNA per reaction (e.g., Cao et al. 2007). In contrast, qPCR systems generally exhibit greater sensitivity, with reported detection limits in the low femtogram range, such as 4 fg (Sundelin et al. 2010) and 10 fg (Cao et al. 2014) per reaction. BLAST analysis of all available *P. brassicae* genomes in the NCBI database indicated that each genome contains no more than four copies of the rDNA target. For example, four copies were identified in strain 3A, the only fully sequenced strain available, whereas fewer than four copies were detected in the remaining 51 partially sequenced strains (https://www.ncbi.nlm.nih.gov/datasets/genome/?taxon=37360). The genome size of strain 3A is 25.3 Mb, corresponding to approximately 26 fg of DNA per genome (https://nebiocalculator.neb.com). Because a successful qPCR reaction requires a minimum of three copies of the target sequence (Forootan et al. 2017), the theoretical lower limit of qPCR detection would be at least 9.75 fg of *P. brassicae* DNA per reaction (3/8 × 26 = 9.75). Therefore, caution is warranted when selecting PCR or qPCR systems for clubroot detection if the reported per-reaction detection limit is below 10 fg of DNA.

In addition to the potential inaccuracy of the reported detection limits discussed above, there is a further reason why these values were not used as the metric for comparing the sensitivity of different systems. Substantial inter-laboratory variation exists in DNA extraction efficiency and DNA quantification, particularly for soil-derived samples (Yang et al. 2021). To minimize these sources of variability, this study used synthetic gBlock DNA to compare the sensitivity of the three selected probe-based qPCR systems independently of their previously reported detection limits. The use of gBlock DNA bypassed DNA extraction and concentration measurement steps, enabling a more accurate and direct comparison of sensitivity among the qPCR systems.

Most PCR and qPCR systems have been reported to detect *P. brassicae* in soil samples containing as few as 1 × 10^3^ resting spores per g soil (Faggian et al. 1999; Cao et al. 2007; Rennie et al. 2011; Li et al. 2013), with exceptions that the probe-based qPCR of Wallenhammar et al. (2012) had a detection limit of 3 × 10^3^ spores per g soil and the SYBR Green-based qPCR system of Xing et al. (2021) achieved a lower limit of 1 × 10^2^ spores per g of soil.

Our results showed that the probe-based qPCR system developed by Wallenhammar et al. (2012) could detect *P. brassicae* in soil samples with spore concentrations as low as 1 × 10^3^ spores per g soil. This detection limit is comparable to those reported for other PCR and qPCR systems and is slightly lower than that originally reported by Wallenhammar et al. (2012). For most commercial DNA extraction kits, 100 mg of soil is recommended per extraction. The minimum elution volume is typically 50 µL, and 2 µL of eluted DNA is used per qPCR reaction. Consequently, there is a 25-fold reduction in DNA quantity between the original soil sample used for extraction and the DNA template added to each qPCR reaction. If a 100 mg soil subsample were taken from soil containing 100 spores per g soil, only 10 spores would be subjected to DNA extraction, and DNA equivalent to 0.4 spores would be included in each qPCR reaction. DNA from 0.4 spores would contain, at most, 0.4 × 4 × 2 = 3.2 copies of the target sequence. In practice, obtaining 3.2 copies for each reaction is almost impossible. This is because the efficiency of DNA extraction from soil is generally low and Poisson distribution also causes significant variation on sampling (Yang et al. 2021). Therefore, it would be practically impossible for a qPCR assay to generate a detectable signal from soil containing 100 spores per g soil. Thus, we are confident in concluding that the qPCR system developed by Wallenhammar et al. (2012) represents the most sensitive assay currently available for clubroot detection, when DNA extraction and reaction conditions are comparable to those described in this study.

The sensitivity of the qPCR system developed by Wallenhammar et al. (2012) was further validated on DNA of naturally infested soil samples collected from multiple locations across Alberta. The Cq values obtained from samples collected in Sturgeon County and Athabasca County were 33.4 and 35.8, respectively (Table 1). These values represent the threshold resting spore concentrations in soil capable of inducing clubroot gall formation. Based on the standard curve equation shown in Fig. 2b, the corresponding resting spore concentrations in the original soil samples were estimated to be approximately 2,000 and 1,200 spores per g soil, respectively. These results are consistent with conclusions from a previous study indicating that 1,000 spores per mL soil represents a critical threshold for the appearance of visible clubroot symptoms (Zahr et al. 2022). Collectively, these findings confirm that the Wallenhammar et al. (2012) qPCR system reliably predicts the risk of clubroot development under field conditions.

In summary, although several qPCR systems are available for clubroot detection, our experimental results indicate that the probe-based qPCR system developed by Wallenhammar et al. (2012) is currently the best option. This system demonstrates specificity to *P. brassicae* comparable to that of other systems, while providing superior sensitivity. Adoption of a single, optimized qPCR system would improve the consistency of detection results and facilitate the comparability and interchangeability of data among different diagnostic laboratories. This comparability opens the possibility for more accurate interprovincial evaluations and could be a step toward improved consistency and accuracy as part of a quality-controlled standard method for clubroot resting spore detection and quantification.

## Notes

### Competing Interest Statement

The authors have declared no competing interest.

